# Double-stranded DNA viruses may serve as vectors for horizontal transfer of intron-generating transposons

**DOI:** 10.1101/2025.03.18.643946

**Authors:** Landen Gozashti, Russell Corbett-Detig

## Abstract

Specialized transposable elements capable of generating introns, termed introners, are one of the major drivers of intron gain in eukaryotes. Horizontal gene transfer (HGT) is thought to play an important role in shaping introner distributions. Viruses could function as vehicles of introner HGT since they often integrate into host genomes and have been implicated in widespread HGT in eukaryotes. We annotated integrated viral elements in diverse dinoflagellate genomes with active introners and queried viral elements for introner sequences. We find that 25% of viral elements contain introners. The vast majority of viral elements represent maverick-polinton-like double-stranded DNA (dsDNA) viruses as well as giant dsDNA viruses. By querying a previously annotated set of maverick-polinton-like proviruses, we show that introners populate full-length elements with machinery required for transposition as well as viral infection. Introners in the vast majority of viral elements are younger than or similar in age to others in their host genome, suggesting that most viral elements acquired introners after integration. However, a subset of viral elements show the opposite pattern wherein viral introners are significantly older than other introners, possibly consistent with virus-to-host horizontal transfer. Together, our results suggest that dsDNA viruses may serve as vectors for HGT of introners between individuals and species, resulting in the introduction of intron-generating transposons to new lineages.

## Main text

### Introduction

Spliceosomal introns are noncoding elements that interrupt eukaryotic genes and must be removed by the spliceosome before translation. Introns are near-ubiquitous components of eukaryotic genomes and serve important functions (reviewed in [1–3]) but have poorly understood origins. Introners, transposable elements capable of generating new introns upon insertion into exons, are a major source of ongoing intron gain in diverse eukaryotes [4–9]. Horizontal gene transfer (HGT) is thought to play an important role in shaping introner distributions across taxa, and recent work has revealed several possible cases of introner HGT between highly divergent lineages (e.g. between dinoflagellates and sponges) [4,10]. Despite evidence of HGT, the molecular mechanisms by which introners move between species remains a mystery.

One mechanism by which introners could move between individuals and species is by hitchhiking on viruses. Viruses are powerful vehicles for horizontal gene transfer and have frequently been implicated in HGT between species, including HGT of other mobile genetic elements [11–14]. The tendency for many viruses and virus-like transposons to integrate into host genomes allows them to capture and mobilize endogenous DNA between hosts [13–16]. Recent work has shown that viruses that are integrated into host genomes, hereafter referred to as viral elements, are common in species with active introners [14,17,18]. In particular, viral elements stemming from double-stranded DNA viruses related to self-synthesizing maverick-polinton DNA transposons have risen to high copy numbers in dinoflagellates in the genuses *Polarella* and *Symbiodinium* [14,17]. *Polarella* and *Symbiodinium* species contain diverse and abundant active introners, raising the question of whether viruses may act as vectors for HGT within and between species.

To address this question, we queried viral elements for introners in four genomes of dinoflagellate isolates with known introners (genera *Polarella* and *Symbiodinium*). We annotated viral elements *de novo* in each genome and also queried a set of full-length proviruses previously annotated in *Polarella* and *Symbiodinium* genomes. We find that introners are present in many viral elements, including elements with intact machinery for viral replication and morphogenesis. Introners in the majority of viral elements are less divergent or similarly divergent than introners in the rest of the host genome, consistent with secondary insertion of introners after viral integration. However, some viral elements show the opposite pattern, in which viral introners are more divergent than expected. This case is more consistent with a model whereby viral elements introduce new intron-generating transposons to their host. Together, our results suggest that dsDNA viruses may serve as vehicles for HGT of introners between individuals and species.

## Results and discussion

### Double-stranded DNA viral elements are common in dinoflagellate genomes

To search for introners in viral elements, we first annotated candidate viral elements in four dinoflagellate genomes known to harbor active introners (genera *Polarella* and *Symbiodinium*; Supplemental Table S1). We chose to survey these isolates since they harbor diverse and abundant active introners [4,10,19], are host to a wide range of viruses and have been shown to possess integrated viral elements with high copy numbers [17]. We used a recently developed deep neural network method to annotate and classify viral elements in each genome [20], revealing a total of 476 viral elements that could be confidently classified. Among these, the majority comprised double-stranded DNA viruses within the kingdom Bamfordvirae (~75% of total classified viruses), consistent with massive invasions of integrating dsDNA viruses recently reported in dinoflagellates [14,17] (**Figure 1A**). Although many viral elements could only be classified at higher taxonomic levels (e.g. kingdom, Bamfordvirae; realm, Duplodnaviria), most elements that could be classified at lower taxonomic levels (class or order) were classified as adintoviruses. Viral elements within this clade are known as maverick-polinton-like elements due to their phylogenetic relationship to self-synthesizing maverick and polinton transposable elements [21]. Most other viral elements with lower-level classifications reside within the families Phycodnaviridae and Tectiviridae. Viruses in the family Phycodnaviridae represent giant viruses which have frequently been implicated in HGT in protists including dinoflagellates [13]. Viral elements classified as tectiviruses probably primarily represent additional misclassified maverick-polinton-like viral elements due to the close evolutionary proximity between tectiviruses and maverick-polinton-like viruses as well as the fact that tectiviruses are only known to infect bacteria [22]. Together, these results suggest that while diverse dsDNA viral elements likely occupy dinoflagellate genomes, maverick-polinton-like viral elements and giant virus related viral elements are particularly common.

**Figure 1:**
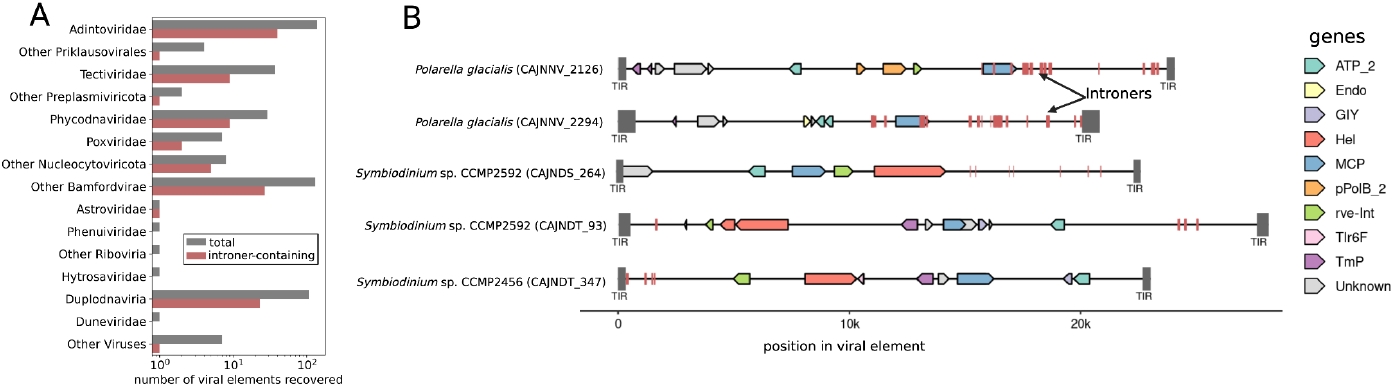
(**A**) The total number of viral elements recovered from considered dinoflagellate genomes using genomad [20]. (**B**) Examples of introner-containing full-length maverick-polinton-like viral elements. Viral genes, introners and flanking terminal inverted repeats (TIRs) are annotated with arrows, red boxes and gray boxes, respectively. Viral gene abbreviations: ATP_2=packaging ATPase 2, Endo=endonuclease, GIY=GIY-YIG endonuclease, Hel=viral helicase, MCP=Major capsid protein, pPolB_2=protein-primed B-family DNA polymerase 2, rve-Int=retroviral integrase, Tlr6f=Tlr 6F-like protein, TmP=tape-measure protein.

### Introners populate viral elements in diverse dinoflagellates

We next surveyed candidate viral elements for introners using curated introner consensus sequences mined from each considered genome. Introners are present in ~25% of classified viral elements and have colonized at least one viral element across all viral lineages with at least two representative elements (**Figure 1A**). Furthermore, many viral elements contain introners at high copy numbers. Indeed, a single viral element can harbor tens of introners spanning multiple unrelated introner families (**Supplemental Table S2**). Together, these results demonstrate that introners are common in viral elements.

Due to the challenges of accurately annotating full-length proviruses on genomic scales, we also surveyed a previously curated set of proviral elements in dinoflagellate genomes [17]. We identified many examples of integrated full-length maverick-polinton-like dsDNA proviruses harboring introners. Several of these elements are flanked by terminal inverted repeats and possess the machinery required for self-synthesising DNA transposition. Indeed many encode a DNA polymerase or viral helicase and an integrase as well as additional cargo previously observed in self-synthesising transposons [16,23] (**Figure 1B**). However, introner-harboring proviruses also possess machinery required for viral morphogenesis such as genes encoding capsid proteins, suggesting that they could still function as infectious viruses [23] (**Figure 1B**). Maverick-polinton-like dsDNA elements have been shown to serve as powerful vectors for HGT, including HGT of other mobile genetic elements [15,24]. The presence of introners in many full-length integrated maverick-polinton-like dsDNA elements highlights the potential for these elements to spread introners within and between species.

### Molecular evidence of secondary insertion into viral elements as well as virus-mediated introduction

Two non-mutually-exclusive hypotheses could explain the presence of introners within viral elements. First, introners already present in a host could insert into a viral element post-integration (secondary insertion hypothesis). Second, an exogenous viral element could introduce introners into a genome upon integration, which did not previously occur in the respective host’s genome (viral introduction hypothesis). One expectation for the viral introduction hypothesis is that introners within a viral element should be more divergent from each other than introners in the host genome. Indeed, if a viral element was the initial source of introners in the host genome, we would expect that viral introners are older than those in the host genome on average. The secondary insertion hypothesis predicts the opposite trend, in which introners within a particular viral element are younger or similar in age to introners in the host genome.

We performed permutation tests in which we computed the mean pairwise divergence between introners in a viral element and compared to expectations from randomly resampled introners in the respective host genome across 1000 permutations. When viral elements harbored multiple introner families, we performed independent tests for each introner family present. We omitted introner copies primarily composed of low complexity regions and only considered cases in which a viral element harbored at least 5 introner copies. After applying these filters, we tested 197 viral element-introner pairs. The vast majority of these (~90%) showed patterns more consistent with the secondary insertion hypothesis in which the mean pairwise divergence of viral introners was less than or did not significantly deviate from expectations. This suggests that secondary insertion of introners into integrated viruses is likely much more common than virus-mediated introner introduction. However, we also found a 10% subset of viral element-introner pairs that showed patterns more consistent with the viral introduction hypothesis. In these cases, introners in viral elements were significantly more divergent from each other than expected from introners in the host genome, suggesting that introner insertions within the viral element might predate other introners in the genome. These viruses and introners represent interesting candidates for downstream scrutiny. Nonetheless, we note this observation could also be caused by variation in mutation rates across the genome [25] or, in cases where the viral element is still actively mobilizing, higher mutation rates in the viral element itself [26]. Overall, these results suggest that the vast majority of introner insertions in viral elements occurred after viral integration.

### Do viral elements serve as vehicles for introner transmission?

Introner-containing species show patchy taxonomic distributions and are enriched for aquatic unicellulars and ascomycete fungi –lineages which generally experience high rates of HGT as a result of germline accessibility and ecology as well as a myriad of other factors [4,27–29]. This circumstantial evidence as well as more recent molecular evidence of HGT of introners between divergent species suggests that HGT plays an important role in shaping introner distributions and, consequently, the distribution of intron gain across taxa [4,10]. Viruses are obvious candidates for vehicles of introner HGT since they often integrate into host genomes and have been implicated in widespread HGT [11–13]. Here, we show that introners are common in dsDNA viral elements in dinoflagellate genomes where both have been tremendously successful [17,19]. Previous observations of introners in genes of viral origin hinted at the possibility of virus-mediated HGT [10]. Our results here shed additional light on this hypothesis and constitute the first observation of introners in putative viral elements as well as full-length proviruses. The majority of introners in viral elements show evidence that they inserted following viral integration. Nonetheless, integrated viral elements in dinoflagellates and other species can represent viruses in a stage of latency, which can become infectious at a later time upon reactivation [14,30,31]. Notably, this strategy is commonly employed by dsDNA viruses closely related to the introner harboring viral elements identified here [14,30,32,33]. Thus, it remains entirely plausible that viral elements, which acquire introners post-integration, may still function as active viruses and serve as vehicles for HGT of introners between species.

Several full-length maverick-polinton-like proviruses harbor introners and also retain genes required for viral morphogenesis, which are usually lost from endogenous viruses or self-synthesizing transposons, providing further evidence that many of these may indeed constitute latent viruses and/or virophages [17,23,34]. Introners have also colonized viral elements originating from giant dsDNA viruses (Phycodnaviridae and other viruses in the phylum Nucleocytoviricota), which are known to frequently acquire host endogenous DNA and facilitate HGT between a diverse breadth of eukaryotic hosts [13]. Indeed, a recent survey found that giant dsDNA viruses account for >80% of detectable virus-eukaryote genetic exchanges [13]. In light of the presence and abundance of introners in dsDNA viral elements that are well-established vectors for HGT, we postulate that virus-mediated HGT of introners likely occurs. However, comprehensive searches for introners in exogenous viral genomes have not yet yielded any promising results. Our ability to explore dsDNA viruses and dsDNA viral elements in introner-containing species on genomic scales has been limited by a lack of comprehensive methods for viral element identification as well as a dearth of high-quality protist and protist-infecting virus genomes [35–37]. Nonetheless, these findings underscore the potential role of dsDNA viruses in shaping the evolutionary landscape of eukaryotic genomes by facilitating the horizontal transfer of intron-generating transposons across diverse lineages.

## Methods

### Retrieving relevant genomic data

We retrieved all considered dinoflagellate genome assemblies from NCBI. Accessions are listed in Supplemental Table S1. Introner consensus sequences for each genome were previously mined and curated [10], and are publicly available at https://github.com/lgozasht/Introner-elements. Previously identified maverick-polinton-like viral elements were retrieved from [17].

### *De novo* viral element annotation

We used geNomad to annotate viral elements *de novo* in each genome [20]. geNomad is an annotation tool that combines information from gene content and a deep neural network to identify and classify viral elements in assembled nucleotide sequences with high precision and accuracy [20]. We ran genomad end-to-end with default parameters on each genome. GeNomad results for each genome are summarized in Supplemental Tables S3-6. Many candidate mobile elements discovered with GeNomad remained unclassified since they did not possess any viral hallmarks (**Supplemental Tables S3-6**). We excluded these elements from downstream analyses.

### Annotating introners in viral elements

To search of introners in viral elements, we first performed blastn [38] searches with parameters-task blastn -evalue 0.1 -outfmt “6 qseqid sseqid qcovs qcovhsp pident length mismatch gapopen qstart qend sstart send evalue bitscore stitle” to query viral elements for introner sequences corresponding to each respective genome. We filtered these results for hits with query coverage ≥90 and alignment length >40. Then, we used bedtools to sort and merge overlapping hits in each viral element. We did this for both viral elements mined de novo in this study and a set of full-length maverick-polinton-like proviruses previously mined in [17]. GeNomad results likely include the viral elements mined in [17] in addition to a more exhaustive set of other diverse viruses. Thus, we used GeNomad results for all downstream calculations and analyses, with the exception of figure 1A. Identifying and annotating full-length intact proviruses usually requires additional curation. Thus, rather than processing viral elements in our GeNomad results, we opted to use the set of maverick-polinton elements previously identified in [17] in order to confirm the presence of introners in intact proviruses. We used gggenomes [39] to virtualize gene and introner annotations in proviruses.

### Exploring molecular evidence introner insertion ages in viral elements compared to genome-wide expectations

Assuming that introners in viral elements and introners in eukaryotic host genomes experience equal rates of molecular evolution, if introner activity in a viral element predated viral integration, introners in that viral element should be more divergent from each other than introners in the rest of the host genome (viral introduction hypothesis). Any case in which introners in a viral element are similarly divergent from each other or less divergent from each other than genome-wide expectations would support the opposite scenario, in which introners inserted into a viral element post-integration (secondary insertion hypothesis). To test between these two hypotheses, we performed permutation tests in which we compared the mean pairwise divergence for introners in each viral element to expectations from randomly resampled introners in the corresponding host’s genome. We performed these tests across 1000 permutations and only considered cases in which a viral element harbored at least 5 copies from the same introner family. Prior to performing these tests, we also used DUST [40] to filter all introners primarily composed of low complexity regions. Multiple sequence alignments were generated using MAFFT [41] and mean pairwise divergence was computed with BioPython [42]. Permutation test results are reported in Supplemental Table S7.

## Supplemental Table Captions

**Supplemental Table S1:** Assembly accessions for dinoflagellate genomes considered in this study.

**Supplemental Table S2:** Number of introners identified for each introner family in each viral element across all considered genomes.

**Supplemental Table S3:** GeNomad results for *Polarella glacialis* CCMP2088.

**Supplemental Table S4:** GeNomad results for *Polarella glacialis* CCMP1383.

**Supplemental Table S5:** GeNomad results for *Symbiodinium sp*. CCMP2592.

**Supplemental Table S6:** GeNomad results for *Symbiodinium sp*. CCMP2456.

**Supplemental Table S7:** Permutation test results.

## Declarations

## Ethics approval and consent to participate

Not applicable

## Consent for publication

Not applicable

## Availability of data and materials

Genomes analysed in the current study are available from NCBI. Corresponding NCBI accessions are presented in Supplemental Table S1. Introner consensus sequences are publicly available at https://github.com/lgozasht/Introner-elements. Previously identified maverick-polinton-like viral elements were identified in [17] and can be retrieved from https://doi.org/10.6084/m9.figshare.21581355.v3.

## Author contributions

L.G. and R.C.-D. conceived and designed the research. L.G. performed all data analyses and data visualization. Both authors wrote the manuscript. Both authors edited and contributed to the manuscript revision.

## Acknowledgements

The authors thank Chris Condon and Peter Sudmant for helpful discussions and feedback on this manuscript.

